# Consistent changes in the intestinal microbiota of Atlantic salmon fed insect meal diets

**DOI:** 10.1101/2021.07.04.451034

**Authors:** Yanxian Li, Karina Gajardo, Alexander Jaramillo-Torres, Trond M. Kortner, Åshild Krogdahl

## Abstract

Being part of fish’s natural diets, insects have become a realistic, sustainable feed ingredient for aquaculture. While nutritional values of insects have been extensively studied in various fish species, their impact on the fish microbiota remains to be fully explored. In an 8-week freshwater feeding trial, Atlantic salmon (*Salmo salar*) were fed either a commercially relevant reference diet or an insect meal diet wherein black soldier fly (*Hermetia illucens*) larvae meal comprised 60% of total ingredients. Microbiota of digesta and mucosa origin from the proximal and distal intestine were collected and profiled along with feed and water samples. The insect meal diet markedly modulated the salmon intestinal microbiota. Overall, the microbial diversity was lower in the digesta of salmon fed the insect meal diet but higher in the mucosa. A group of bacterial genera, dominated by members of the *Bacillaceae* family, was enriched in salmon fed the insect meal diet, which confirms our previous findings in a seawater feeding trial. We also found that microbiota in the intestine closely resembled that of the feeds but was distinct from the water microbiota. Notably, bacterial genera associated with the diet effects were present in the feeds as well. In conclusion, our results show consistent changes in the intestinal microbiota of Atlantic salmon fed diets containing black soldier fly larvae meal.

## Introduction

The global population is projected to reach 9.7 billion in 2050 (1), requiring an increase in the food supply by 25-70% (2). To fulfill this demand, the food production sector must minimize resource input and maximize nutritional outputs for human consumptions. Atlantic salmon, *Salmo salar*, is the most produced marine fish species and one of the most economically important farmed fish worldwide (3). Human-edible plant feedstuffs are the main ingredients used in modern salmon feeds (~70%) (4). To secure sustainable developments, salmon farming needs to decrease its dependency on human-edible feedstuffs and incorporate unexploited feed resources in its raw material repertoire. So far, possible candidates include insects (5), macroalgae (6), and single-cell organisms such as bacteria, yeasts, and microalgae (7). In terms of sustainability, insects are a promising candidate. They possess a remarkable capacity to upgrade low-quality organic materials, require minimal water and cultivable land, and emit little greenhouse gases (8). One insect species with the potential as an alternative protein source for salmon aquaculture is the black soldier fly (*Hermetia illucens*), produced at an industrial scale for its good nutritional value (9). Feed conversion ratio, growth performance, and fish health are primary concerns when evaluating the performance of alternative feed ingredients. While the nutritional value of black soldier fly larvae meal has been extensively evaluated in various fish species, including Atlantic salmon (10–16), its influence on fish health remains largely unexplored.

The intestine is the first organ exposed to the diet and of pivotal importance for the growth, development, and protection against pathogens. A well-functioning, healthy intestine is the key to convert feed into fish biomass efficiently. It is now well established that the intestinal microbiota is, in various ways, closely connected to intestinal function and health (17–21). Diet is arguably one of the most important environmental factors shaping intestinal microbiota (22–24). Different dietary components may selectively induce compositional and functional alterations of the intestinal microbiota, which in turn could inflict important implications on the host health and disease resistance (19,24–26).

Characterizing the response of intestinal microbiota to dietary shifts and its associations with host responses is a critical step towards identifying key microbial clades for promoting fish health and welfare. The main aims of the work presented herein were (i) to compare intestinal microbiota of Atlantic salmon fed a commercially relevant reference diet and an insect meal-based test diet, and (ii) to identify potential associations between intestinal microbial clades and host responses. This work was part of a larger study consisting of a freshwater and a seawater feeding trial. The present work reports the intestinal microbiota in freshwater Atlantic salmon fed an insect meal diet containing 60% black soldier fly larvae meal for 8 weeks.

## Materials and methods

### Experimental fish, diet and sampling

An 8-week fresh-water feeding trial was conducted at Cargill AquaNutrition experimental facility at Dirdal, Norway, in compliance with the Norwegian Animal Welfare Act 10/07/2009 and the EU Directive on the Protection of Animals used for Scientific Purposes (2010/63/EU). A total of 800 Atlantic salmon with a mean initial body weight of 49 g (1.5 g SEM) were randomly assigned into 8 fiberglass tanks (450 L, 100 fish per tank) supplied with running freshwater. Quadruplicate tanks of fish were fed either a reference diet with a combination of fish meal, soy protein concentrate, and wheat gluten as protein sources, or an insect meal diet wherein 85% of the protein was supplied by black soldier fly larvae meal, replacing most of the fish meal and soy protein concentrate (Table 1). The black soldier fly larvae were grown on feed substrates containing organic waste streams. After eight days of growing, the larvae were harvested and partially defatted before being dried and ground to make the insect meal (Protix Biosystems BV, Dongen, The Netherlands). The insect meal contains about 40% crude protein and 34% crude lipid. The diets were formulated and produced by Cargill (Dirdal, Norway) and stored at −20 °C until use. The fish were fed continuously by automatic disk feeders under a photoperiod regimen of 24 h daylight. Uneaten feeds were collected from tank outlets and registered daily. During the feeding trial, the water temperature was 13.7 ± 0.1 °C, and the dissolved oxygen concentration of the inlet and outlet water was 11.9 ± 1.2 and 8.7 ± 0.5 mg/L, respectively. Further details on the nutritional composition of the insect meal and diets have been reported elsewhere (27,28).

**Table 1.**
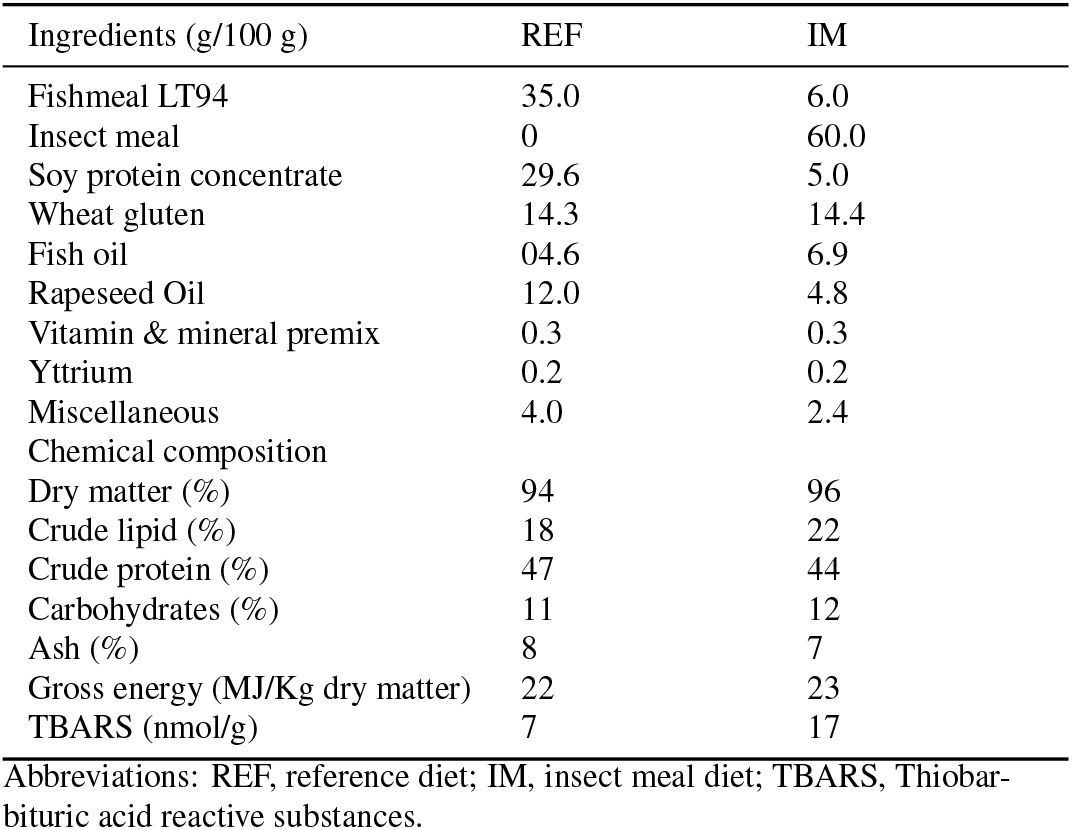
Formulation of the experimental diets.

### Sample collection

At the termination of the feeding trial, 3 fish were randomly taken from each tank (i.e., 12 fish per treatment), anesthetized with tricaine methanesulfonate (MS222^®^; Argent Chemical Laboratories, Redmond, WA, USA), and euthanized by a sharp blow to the head. After cleaning the exterior of each fish with 70% ethanol, the proximal and distal intestine were aseptically removed from the abdominal cavity, placed in sterile Petri dishes, and opened longitudinally. Only fish with digesta along the whole intestine were sampled to ensure that the intestine had been exposed to the diets. The intestinal digesta was gently removed and transferred into a 1.5 mL sterile Eppendorf tube using a spatula and snap-frozen in liquid N_2_ for the profiling of digesta-associated intestinal microbiota. The intestinal tissue was rinsed in sterile phosphate-buffered saline 3 times to remove traces of remaining digesta. After rinsing, the intestinal tissue was cut into 3 pieces for histological evaluation (fixed in 4% phosphate-buffered formaldehyde solution for 24 h and transferred to 70% ethanol for storage), gene expression analysis (preserved in RNAlater solution and stored at −20 °C), and profiling of mucosa-associated intestinal microbiota (snap-frozen in liquid N_2_), respectively. In addition, 300 mL water was taken from each tank, pre-filtered through a 0.8 μm sterile syringe filter (Acrodisc®, Pall Corporation, New York, USA), and vacuum-filtered onto a 0.2 μm sterile nitro-cellulose filter (Nalgene™, Thermo Scientific, USA). The filter containing enriched bacteria was folded, placed into an 8 mL sterile tube, and snap-frozen in liquid N_2_ to profile microbial community in water. The collection of microbiota samples was performed near a gas burner to secure aseptic conditions. Tools were cleaned and decontaminated by 70% ethanol sprays and flaming before the subsequent sampling was carried out. The samples for microbiota profiling were transported in dry ice and stored at −80 °C until DNA extraction.

### DNA extraction

Total DNA was extracted from ~100 mg digesta, mucosa, and feed using the QIAamp DNA Stool Mini Kit (Qiagen, Hilden, Germany) as previously described (29), except that 2 mL prefilled PowerBead tubes (glass beads, 0.1 mm; Cat no. 13118-50, Qiagen) were used for the bead beating. To extract DNA from water samples, the frozen filter was allowed to soften on ice and rolled into a cylinder with the white filter membrane facing outward using two sets of sterile forceps. The filter was then inserted into an 8 mL sterile tube containing the double amount of ASL buffer and glass beads used in the prefilled PowerBead tubes. The tube was secured horizontally to a mixer mill (Retsch GmbH, Germany; model, MM 301) and shaken vigorously at the frequency of 30 Hz for 5 min (2.5 min, pause and invert the tube, 2.5 min). After shaking, the tube was centrifuged at 4000 g for 1 min, and 2.6 mL supernatant was collected and evenly aliquoted into two 1.5 mL Eppendorf tubes. The DNA was extracted from the supernatant aliquots and pooled afterward, following the protocol as previously described (29). For quality control purposes, a companion “blank extraction” sample was added to each batch of sample DNA extraction by omitting the input material, whereas an additional mock sample (ZymoBIOMICS™, Zymo Research, California, USA; catalog no., D6300) was included for each DNA extraction kit as a positive control. The mock consists of 8 bacteria (*Pseudomonas aeruginosa, Escherichia coli, Salmonella enterica, Lactobacillus fermentum, Enterococcus faecalis, Staphylococcus aureus, Listeria monocytogenes*, *Bacillus subtilis*) and 2 yeasts (*Saccharomyces cerevisiae*, *Cryptococcus neoformans*).

### Library preparation and sequencing

The V1-2 hyper-variable regions of the bacterial 16S rRNA gene were amplified using the primer set 27F (5’-AGA GTT TGA TCM TGG CTC AG-3’) and 338R (5’-GCW GCC WCC CGT AGG WGT-3’) (30). The PCR was run in a total reaction volume of 25 μL containing 12.5 μL of Phusion®High-Fidelity PCR Master Mix (Thermo Scientific, CA, USA; catalog no., F531L), 10.5 μL molecular grade H_2_O, 1 μL DNA template, and 0.5 μL of each primer (10 μM). The amplification program was set as follows: initial denaturation at 98 °C for 3 min; 35 cycles of denaturation at 98 °C for 15 s, annealing decreasing from 63 °C to 53 °C in 10 cycles for 30 s followed by 25 cycles at 53 °C for 30 s, and extension at 72 °C for 30 s; followed by a final extension at 72 °C for 10 min. The PCR was run in duplicate incorporating negative PCR controls, which were generated by replacing the template DNA with molecular grade H_2_O. The duplicate PCR products were pooled and examined by a 1.5% agarose gel electrophoresis before cleanup.

The sequencing was carried out on a Miseq platform following the Illumina 16S metagenomic sequencing library preparation protocol (31). Briefly, the PCR products were cleaned using the Agencourt AMPure XP system (Beckman Coulter, Indiana, USA; catalog no., A63881), multiplexed by dual indexing using the Nextera XT Index Kit (Illumina, California, USA; catalog no., FC-131-1096) and purified again using the AMPure beads. After the second clean-up, representative libraries were selected and analyzed using the Agilent DNA 1000 Kit (Agilent Technologies, California, USA; catalog no., 5067-1505) to verify the library size. Cleaned libraries were quantified using the Invitrogen Qubit^TM^ dsDNA HS Assay Kit (Thermo Fisher Scientific, California, USA; catalog no., Q32854), diluted to 4 nM in 10 mM Tris (pH 8.5) and finally pooled in an equal volume. Negative controls with library concentrations lower than 4 nM were pooled in equal volume directly. Due to the low diversity of amplicon library, 15% Illumina generated PhiX control (catalog no., FC-110-3001) was spiked in by combining 510 μL amplicon library with 90 μL PhiX control library. The library was loaded at 6 pM and sequenced using the Miseq Reagent Kit v3 (600-cycle) (Illumina; catalog no., MS-102-3003).

Due to technical challenges in obtaining high-quality PCR products for mucosa samples, the digesta samples were first amplified and sequenced. The PCR conditions for mucosa samples were optimized by diluting the DNA templates (1:5) to reduce the influence of PCR inhibitors. The mucosa samples were then sequenced in a second run together with feed and water samples. To assess potential batch effects between sequencing runs, 8 representative digesta samples were also sequenced in the second run to serve as technical replicates.

### Sequence data processing

The raw sequence data from each run were separately processed by the DADA2 (version 1.18) in R (version 4.0.5) (32) to infer amplicon sequence variants (ASVs) (33). Specifically, the demultiplexed paired-ended reads were trimmed off the primer sequences (first 20 bps of forward reads and first 18 bps of reverse reads), truncated at the position where the median Phred quality score crashed (forward reads at position 260 bp and reverse reads at position 188 bp for the first run; forward reads at position 290 bp and reverse reads at position 248 bp for the second run) and filtered off low-quality reads. After the trimming and filtering, run-specific error rates were estimated, and the ASVs were inferred from each sample independently. The chimeras were removed using the “consensus” method after merging the forward and reverse reads. The resulting feature table and representative sequences from each run were imported into QIIME2 (version 2020.11) (34) and merged. The taxonomy was assigned by a scikit-learn naive Bayes machine-learning classifier (35), which was trained on the SILVA 132 99% OTUs (36) that were trimmed to only include V1-V2 regions of the 16S rRNA gene. Taxa identified as chloroplasts or mitochondria were excluded from the feature table. The feature table was conservatively filtered to remove ASVs that had no phylum-level taxonomic assignments or appeared in only one biological sample. Contaminating ASVs were identified and removed based on two suggested criteria: contaminants are often found in negative controls and inversely correlate with sample DNA concentration (37), which was quantified by qPCR as previously described (38). The ASVs filtered from the feature table were also removed from the representative sequences, which were then clustered into OTUs at 97% similarity using the VSEARCH *de novo* clustering method (39). The resulting OTU table and representative sequences were used for the downstream data analysis. The phylogeny was constructed by inserting the representative sequences into a reference phylogenetic tree built on the SILVA 128 database using SEPP (40). The alpha-diversity indices were computed by rarefying the OTU table at a subsampling depth of 10 345 sequences. To compare beta-diversity, we performed robust Aitchison PCA using the QIIME2 library DEICODE (41), which is a form of Aitchison distance that is robust to high levels of sparsity in the microbiome data via matrix completion. For downstream data visualization and statistical analyses, QIIME2 artifacts were imported into R using the qiime2R package (42) and a phyloseq (43) object was assembled. As the technical replicates showed strong batch effects between the sequencing runs, which could not be effectively removed by existing batch effect correction methods such as RUVSeq (44) and ComBat-seq (45), we performed the down-stream data analysis independently for samples sequenced in different runs.

### Statistics

Differences in the alpha-diversity indices were compared by linear mixed-effects models using the R package afex (46), which runs the lme4 (47) under the hood to fit mixed-effects models. Predictor variables in the models include the fixed effects Diet + Segment + Diet x Segment, and the random effects FishID + Tank. The models were validated by visual inspections of residual diagnostic plots generated by the R package ggResidpanel (48). The statistical significance of fixed predictors was estimated by Type III ANOVA with Kenward-Roger’s approximation (49) of denominator degrees of freedom. When the interaction between the main effects was significant, conditional contrasts for the main effects were made using the R package emmeans (50). To compare differences in the beta-diversity, we performed the PERMANOVA (51) in PRIMER v7 (Primer-E Ltd., Plymouth, UK) using the same predictors included in the linear mixed-effects models. Terms with negative estimates for components of variation were sequentially removed from the model via term pooling, starting with the one showing the smallest mean squares. At each step, the model was reassessed whether more terms needed to be removed or not. Conditional contrasts for the main effects were constructed when their interaction was significant. Monte Carlo *p* values were computed as well when the unique permutations for the terms in the PERMANOVA were small (< 100). The homogeneity of multivariate dispersions among groups was visually assessed with boxplots and was formally tested by the permutation test, PERMDISP (52), using the R package vegan (53). Per-feature tests for the association between specific microbial clade and sample metadata were done using the R package MaAsLin2 (version 1.4.0) (54). The feature table was collapsed at the genus level and bacterial taxa of low prevalence (present in < 25% of samples) were excluded before running the association analysis. Predictor variables included in the association testing are fixed factors Diet + Segment + *foxp3* (qPCR) + *plin2* (qPCR), and the random effects FishID + Tank. Multiple comparisons were adjusted by the Holm (55) or Benjamini-Hochberg (56) method where applicable. Differences were regarded as significant for *p* < 0.05 or FDR-corrected *q* < 0.1.

## Results

To aid readers in interpreting the data from this feeding trial, results on feed utilization, growth performance, intestinal histomorphology, and gene expression, which have been reported elsewhere (27,57), are summarized as the following. In brief, there was little evidence that the insect meal diet negatively affected salmon’s feed utilization or growth performance. Histopathological examination showed excessive accumulation of lipid (steatosis) in the proximal intestine in both diet groups, but it was less severe in salmon fed the insect meal diet. The expression of the lipid droplet marker gene, *plin2*, supported these histological findings. Immune and barrier-function gene expression profiles were generally not affected by diet. However, salmon fed the insect meal diet showed increased expression of genes indicative of immune tolerance (*foxp3*), stress response (*hsp70*), and detoxification activity (*cpy1a1*).

### Taxonomic analysis

All the bacterial species in the mock were correctly identified at the genus level with *E. faecalis*, *L. fermentum, L. monocytogenes*, and *S. aureus* further being assigned at the species level (Figure S1). At the genus level, the average Pearson’s *r* was 0.58 for the correlation between expected and observed mock composition. The exact sequence, relative abundance, and taxonomy of contaminating features identified in the negative control samples are available in Table S1. The primary contaminating features, in descending order according to the mean relative abundance, were classified as *Pseudomonas*, *Halomonas*, *Shewanella algae*, *Undibacterium*, *Bradyrhizobium*, *Chitinophagaceae*, *Ralstonia, Sediminibacterium, Curvibacter, Afipia*, and *Cutibacterium*.

The top 10 most abundant bacterial genera across all the samples are shown in Figure 1. The microbiota in the digesta collected from the two intestinal segments of the salmon fed the reference diet appeared homogenous, but more heterogeneous in the sampled mucosa. Dominant genera in the reference diet group included *Lactobacillus*, unclassified *Peptostreptococcaceae*, and *Peptostreptococcus*. The microbiota in salmon fed the insect meal diet differed greatly from that of the reference diet fed fish, but the difference between the results of the digesta and mucosa appeared less than for fish fed the reference diet. Dominant genera in the insect meal diet group included unclassified *Bacillaceae*, *Bacillus*, *Corynebacterium 1, Enterococcus, Oceanobacillus*, and *Ornithinibacillus*. The microbiota in the intestine closely resembled that of the feed but was distinct from the water microbiota. In agreement with this, we found that the OTU overlap between the intestine and feed was much higher than that between the intestine and water (Figure 2).

**Fig. 1.**
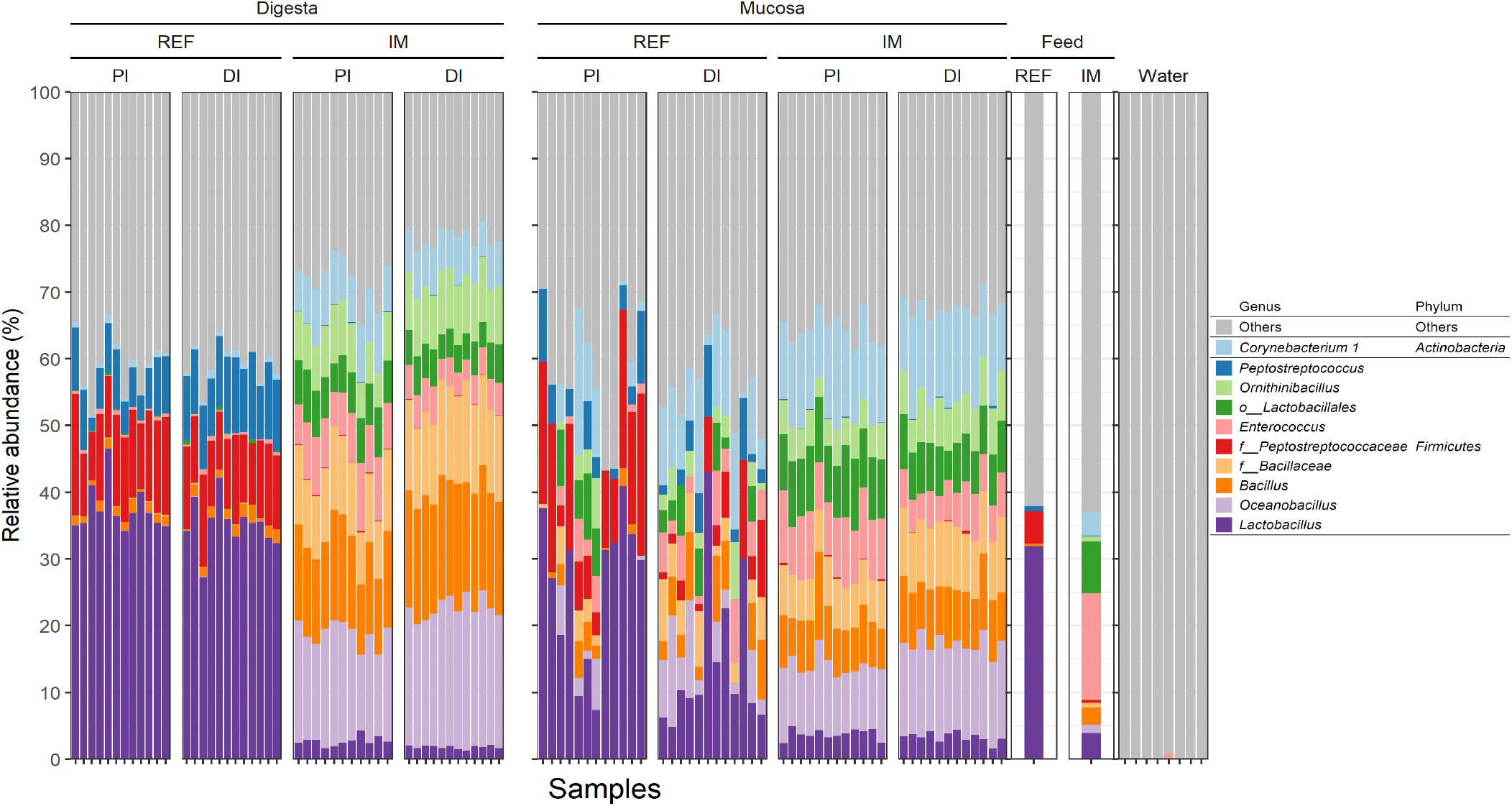
Consistent changes in the taxonomic composition of intestinal microbiota from salmon fed the insect meal diet. Note that feed microbiota shows close resemblance to that observed in the intestine whereas water microbiota is very distinct from the intestinal microbiota. Only the top 10 most abundant bacterial genera are displayed in the plot whereas the other taxa are shown as “Others”. Taxa not assigned at the genus level are prepended with letters indicating whether the taxonomic assignment was made at the order (o_) or family (f_) level. Abbreviations: REF, reference diet; IM, insect meal diet; PI, proximal intestine; DI, distal intestine.

**Fig. 2.**
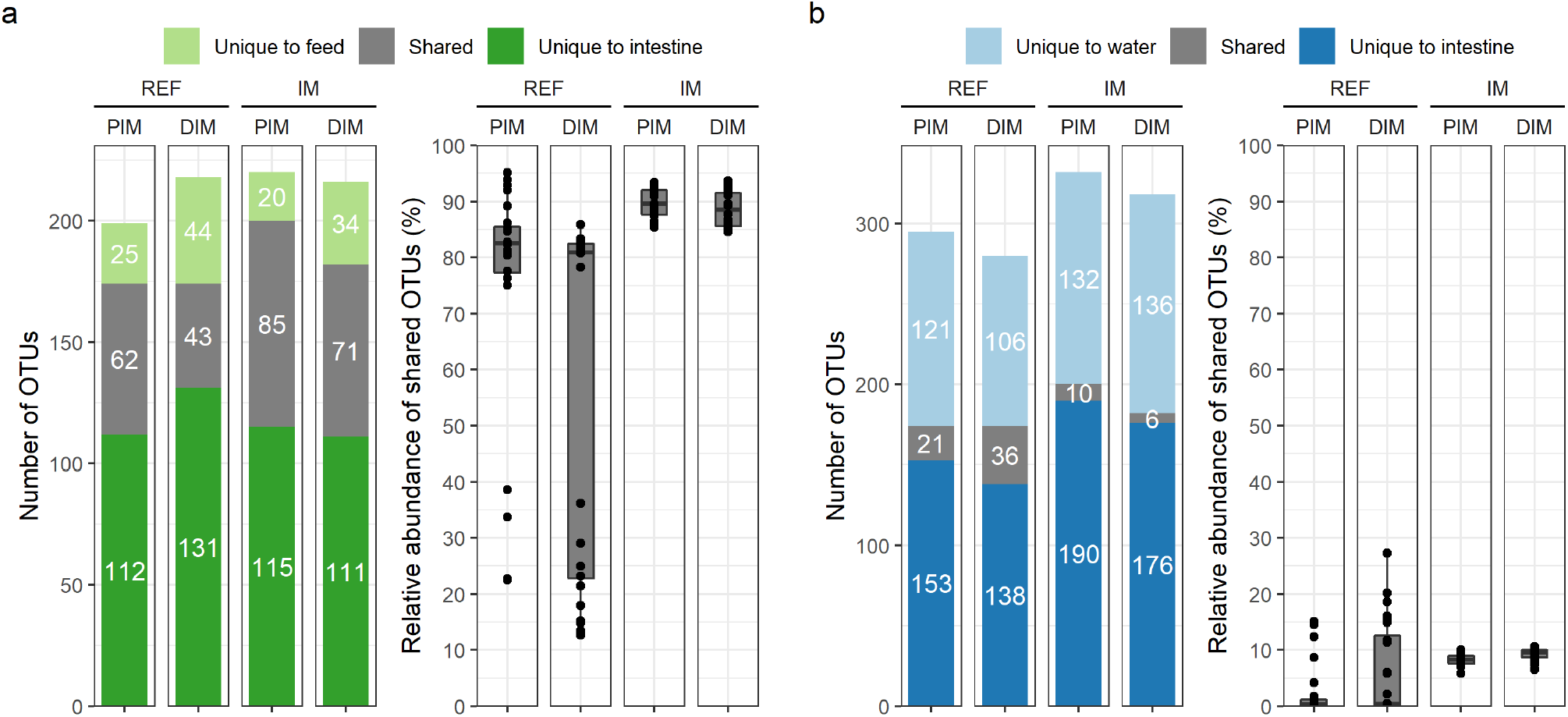
Higher microbial overlap between the intestine and feeds (a) than that between the intestine and water (b). In each panel, the number of shared OTUs is shown on the left whereas the relative abundance of shared OTUs in the intestinal mucosa is shown on the right. To reduce the influence of rare OTUs and differences in sequencing depth, only OTUs with a minimum relative abundance of 0.05% were considered as present in a sample. Abbreviations: REF, reference diet; IM, insect meal diet; PIM, proximal intestine mucosa; DIM, distal intestine mucosa.

### Core microbiota

In total, 45 and 60 OTUs were identified as core microbiota (present in at least 50% of the samples) in salmon fed the reference and insect meal diet, respectively (Figure 3). Five core OTUs were shared between the diets, classified as *Bacillus*, *Globicatella*, *Kurthia*, *Lactobacillus*, and *Ureibacillus*. Primary core OTUs in salmon fed the reference diet comprised *Peptostreptococcaceae* (1 OTU), *Peptostreptococcus* (1 OTU), and lactic acid bacteria including *Lactobacillus* (13 OTUs), *Weissella* (3 OTUs), *Vagococcus* (3 OTUs), *Lactococcus* (1 OTU), *Leuconostoc* (1 OTU), *Pediococcus* (1 OTU) and *Streptococcus* (1 OTU). In contrast, primary core OTUs in salmon fed the insect meal diet comprised *Bacillus* (12 OTUs), *Enterococcus* (7 OTUs), *Corynebacterium 1* (4 OTUs), *Lysinibacillus* (3 OTUs), *Lactobacillus* (3 OTUs), *Actinomyces* (3 OTUs), *Oceanobacillus* (2 OTUs), *Bacillaceae* (2 OTUs), *Brevibacterium* (2 OTUs), *Microbacterium* (2 OTUs), *Ornithinibacillus* (1 OTU) and *RsaHF231*(1 OTU).

**Fig. 3.**
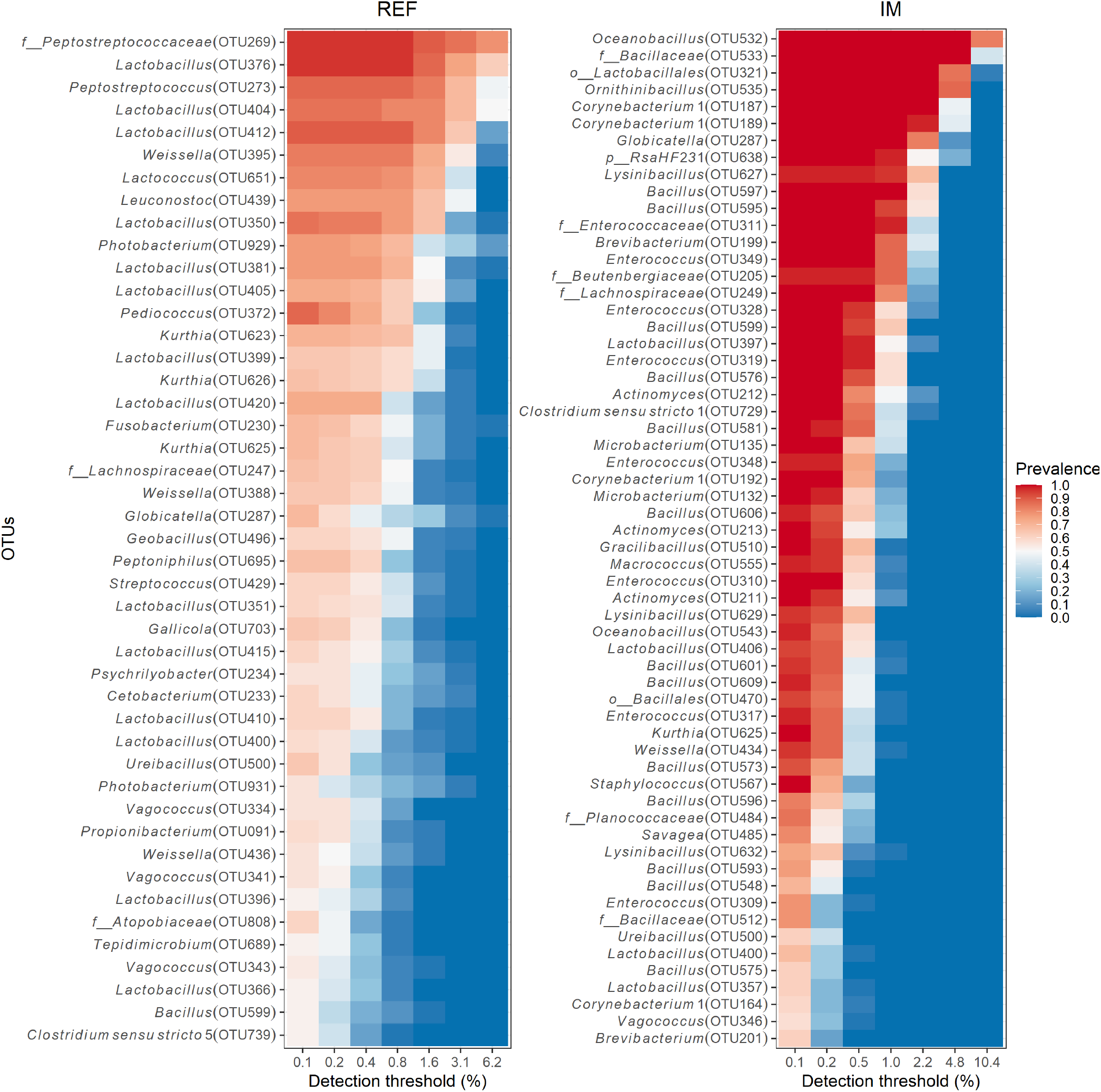
Heatmaps showing the prevalence of core OTUs at different detection thresholds in salmon fed the reference (REF) or insect meal (IM) diet. The core OTUs were computed and visualized by the R package microbiome (58), with a minimum detection threshold of 0.1% and a minimal prevalence threshold of 50%. The taxonomy of core OTUs at the genus level is displayed on the y axis. OTUs not assigned at the genus level are prepended with letters indicating whether the taxonomic assignment was made at the phylum (p_), order (o_), or family (f_) level.

### Alpha-diversity

Overall, the diet effects on the alpha-diversity showed opposite results for digesta and mucosa samples when evaluated independently (Figure 4). In the digesta, the insect meal diet reduced microbial diversity compared to the reference diet, whereas in the mucosa the insect meal diet increased diversity.

**Fig. 4.**
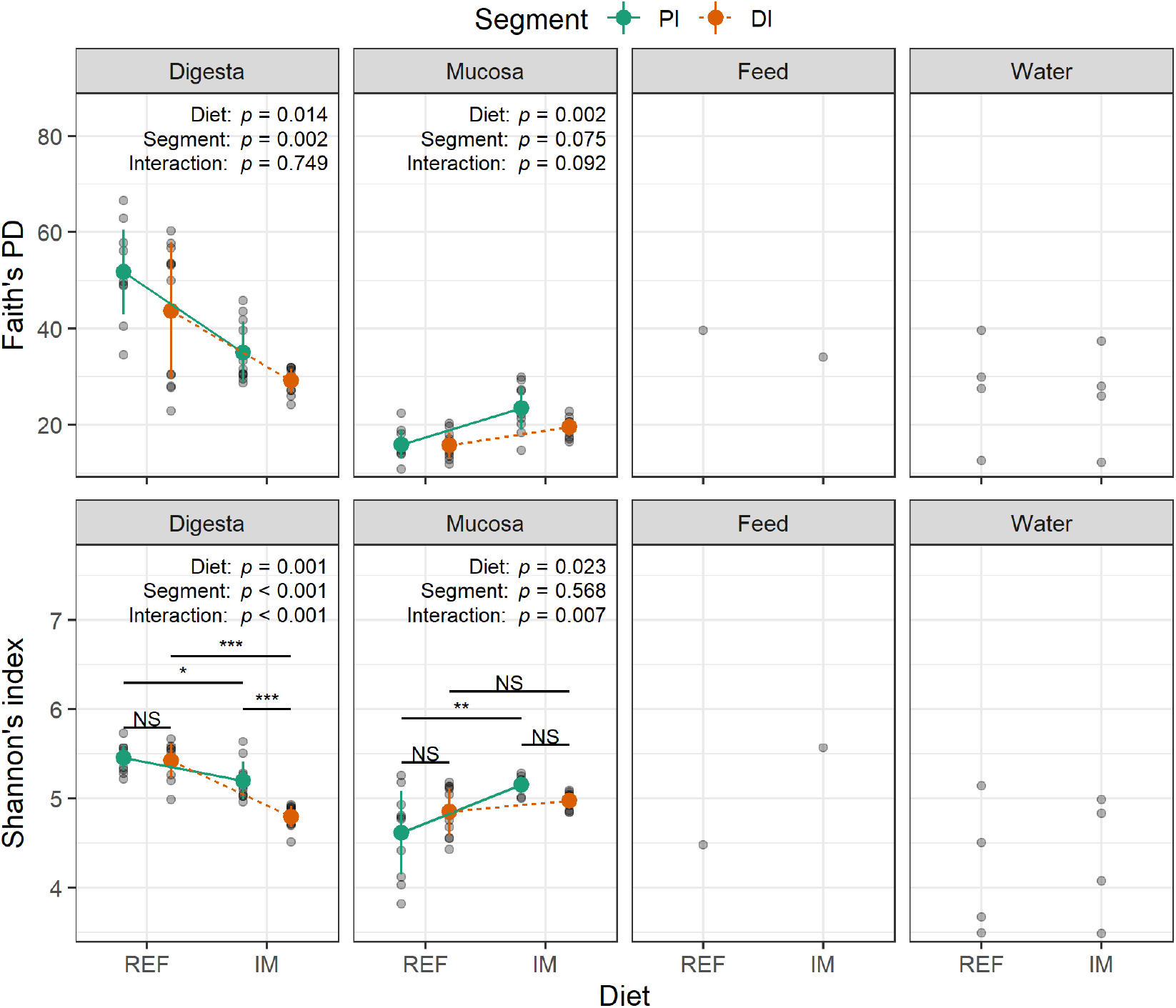
Diet effects on the alpha-diversity are opposite when independently evaluated using digesta or mucosa samples. The error bars denote standard deviations of the means. The *p* values of the main effects and their interaction are displayed on the top of each subplot. The asterisks denote statistically significant differences in the post-hoc conditional contrasts (*, *p* < 0.05; **, *p* < 0.01; ***, *p* < 0.001). Abbreviations: REF, reference diet; IM, insect meal diet; PI, proximal intestine; DI, distal intestine; NS, not significant; PD, phylogenetic diversity.

In the digesta, Faith’s phylogenetic diversity (Faith’s PD) showed a significant diet and intestinal segment effect, and the interaction between these terms was not significant. Faith’s PD was lower in the insect meal diet group than in the reference diet group. Also, it was lower in the distal intestine than in the proximal intestine. Similar results were found for the Shannon index, but the interaction was significant. In both intestinal segments, Shannon’s index was lower in salmon fed the insect meal diet, but the diet effect was stronger in the distal intestine than in the proximal intestine. In contrast, a significant intestinal segment effect was only found in salmon fed the insect meal diet, with the distal intestine showing a lower Shannon’s index.

In the mucosa, Faith’s PD showed a significant diet effect, with no significant difference between the intestinal segments. The salmon fed the insect meal diet showed a higher Faith’s PD in both intestinal segments. A similar effect was indicated by Shannon’s index, but only for the proximal intestine, i.e., the interaction between diet and segment was significant.

The differences between the alpha diversity of water and intestinal mucosal microbiota were not significant (Figure S2).

### Beta-diversity

In the digesta, the PERMANOVA showed a significant diet and intestinal segment effect on the beta-diversity, and the interaction between these terms was significant (Figure 5a; Table 2). The diet effect on the beta-diversity was significant in both intestinal segments, whereas a significant intestinal segment effect was only found in salmon fed the reference diet. The PERMDISP showed that, in both intestinal segments, the multivariate dispersion was higher in the reference diet group than in the insect meal diet group. Differences in the multivariate dispersion between intestinal segments were not significant in both diets (Figure S3a).

**Fig. 5.**
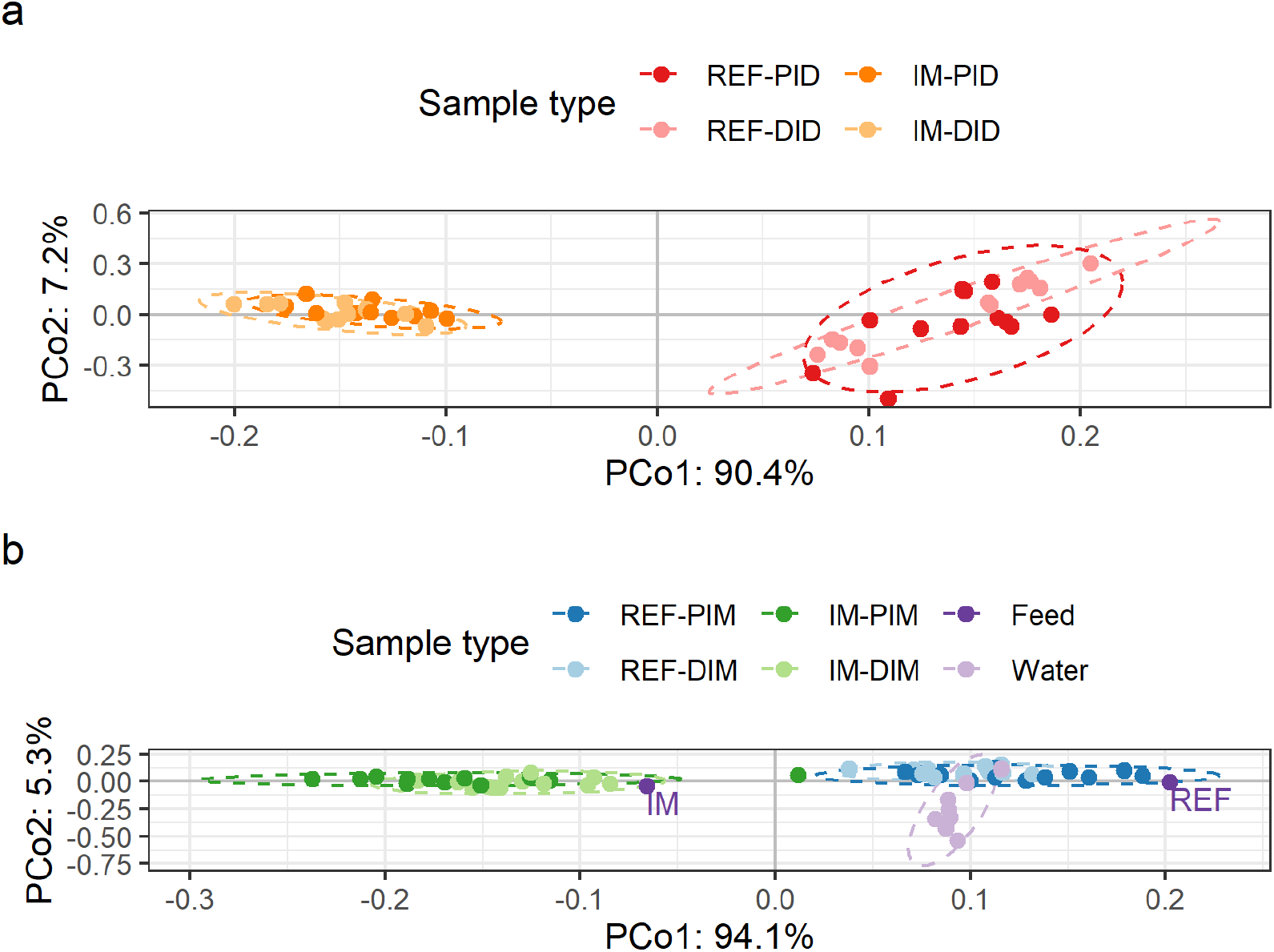
The insect meal diet markedly modulated the salmon intestinal microbiota in both digesta (a) and mucosa (b), irrespective of intestinal segments. The dimensionality reduction was performed using a compositional beta-diversity metric called robust Aitchison PCA and visualized by EMPeror (59). The height-to-width ratio of the PCoA plot was set to reflect the ratio between the corresponding eigenvalues as recommended (60). Abbreviations: REF, reference diet; IM, insect meal diet; PID, proximal intestine digesta; DID, distal intestine digesta; PIM, proximal intestine mucosa; DIM, distal intestine mucosa; PCoA, principal coordinate analysis.

**Table 2.** PERMANOVA and subsequent conditional contrasts.

| Source | Main effects |  | Interaction | Conditional contrasts |  |  |  |
| --- | --- | --- | --- | --- | --- | --- | --- |
|  | Diet | Segment |  | REF-PI VS.<br>IM-PI | REF-DI VS.<br>IM-DI | REF-PI VS.<br>REF-DI | IM-PI VS.<br>IM-DI |
| Digesta | 0.001 | 0.041 | 0.041 | 0.002 | 0.002 | 0.04 | 0.59 |
| Mucosa | 0.001<sup>1</sup> | 0.633 | 0.010 | 0.001<sup>1</sup> | 0.001<sup>1</sup> | NA | NA |

Abbreviations: REF, reference diet; IM, insect meal diet; PI, proximal intestine; DI, distal intestine; NA, not applicable.
<sup>1</sup>Monte Carlo *p* value

In the mucosa, the PERMANOVA showed a significant diet but not a significant intestinal segment effect on the beta-diversity, and the interaction between these terms was significant (Figure 5b; Table 2). The diet effect on the beta-diversity was significant in both intestinal segments, but it was marginally stronger in the proximal intestine than in the distal intestine. The PERMDISP showed that differences in the multivariate dispersion between the diet groups were not significant at the tank or diet level (Figure S3b).

The water microbiota was significantly different from the intestinal mucosal microbiota (*p* = 0.001). The PERMDISP showed that differences in the multivariate dispersion between water and intestinal mucosal samples were not significant (*p* = 0.391).

### Association analysis

Significant associations between sample metadata and bacterial genera in the digesta and mucosa are shown in Figure 6 and Figure 7, respectively. In total, 93 and 36 taxa were associated with the diet effect in the digesta and mucosa, respectively. Collectively, 32 taxa were associated with the diet effect in both digesta and mucosa. Among these taxa, bacterial genera enriched in salmon fed the reference diet consisted of unclassified *Peptostreptococcaceae, Peptostreptococcus, Photobacterium*, and lactic acid bacteria including *Lactobacillus, Lactococcus, Leuconostoc*, *Pediococcus*, and *Streptococcus* (partially illustrated in Figure 6b and Figure 7b). In contrast, bacterial genera enriched in salmon fed the insect meal diet comprised *Actinomyces*, unclassified *Bacillales*, unclassified *Bacillaceae*,*Bacillus*, unclassified *Beutenbergiaceae*, *Brevibacterium, Cellulosimicrobium, Clostridium sensu stricto 1, Corynebacterium 1*, unclassified *Enterococcaceae, Enterococcus*, *Exiguobacterium*, *Globicatella*, *Gracilibacillus*, unclassified *Lactobacillales*, *Lysinibacillus*, *Macrococcus*, *Microbacterium*, *Nosocomiicoccus*, *Oceanobacillus*, *Ornithinibacillus*, *Paenibacillus*, unclassified *Planococcaceae*, and unclassified *RsaHF231* (partially illustrated in Figure 6c and Figure 7c). Regarding associations between bacterial genera and host gene expressions, the relative abundance of *Paenibacillus* and *Streptococcus* in the mucosa showed positive correlations with the expression level of *foxp3*, the master transcription factor of regulatory T-cells, in the intestine (partially illustrated in Figure 7d). Additionally, the relative abundance of unclassified *RsaHF231* in the digesta, and the relative abundance of unclassified *Bacillaceae*, *Corynebacterium 1, Enterococcus*, and *Oceanobacillus* in the mucosa, showed negative correlations with the expression level of *plin2*, a surface marker of lipid droplets, in the intestine (partially illustrated in Figure 7e).

**Fig. 6.**
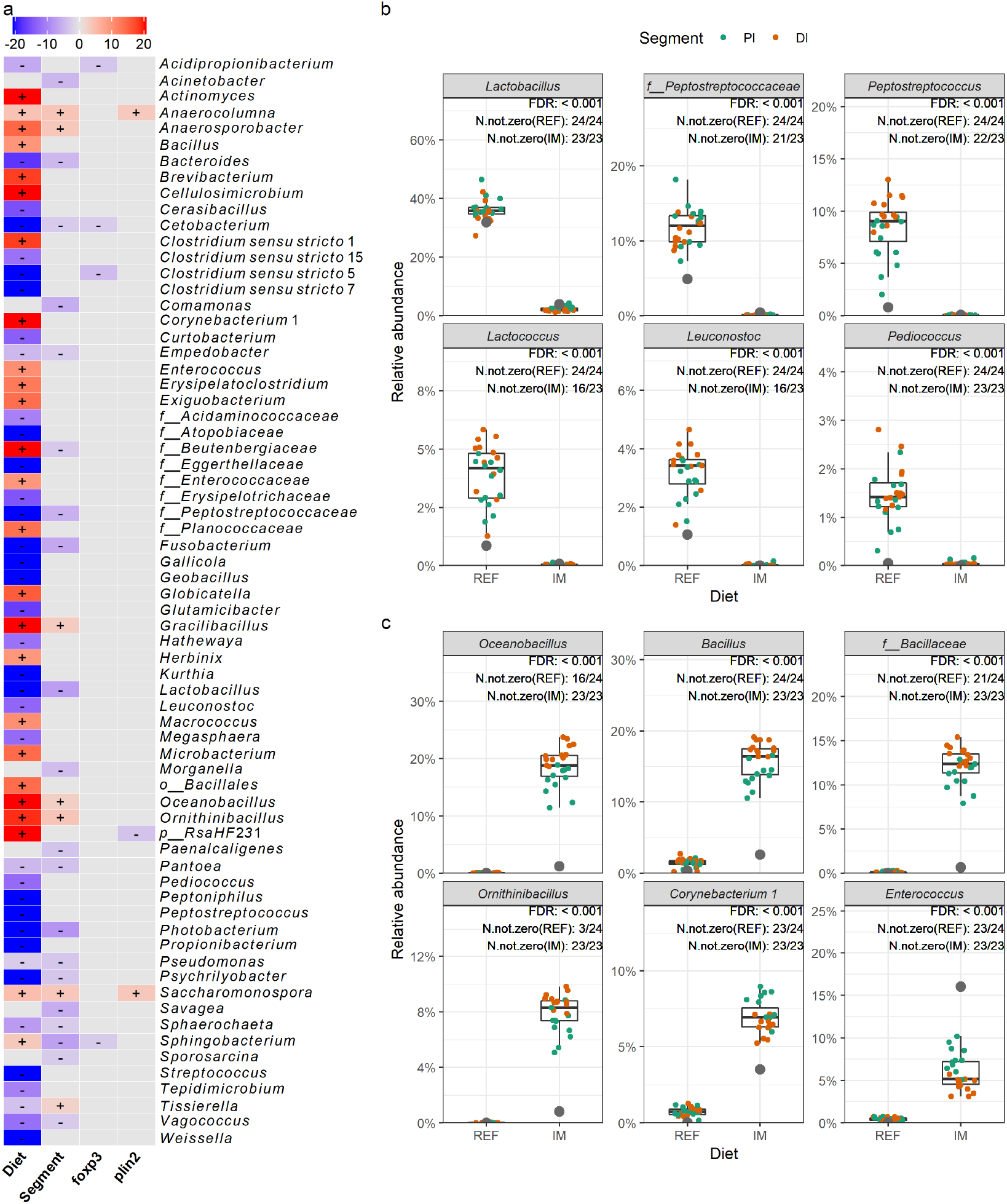
Significant associations between sample metadata and microbial clades in the digesta. (a) Heatmap summarizing significant associations between sample metadata and microbial clades in the digesta. Color key: -log(*q*-value) * sign(coefficient). Cells that denote significant associations are colored in red or blue and overlaid with a plus (+) or minus (-) sign that indicates the direction of association: Diet (+), higher relative abundance in salmon fed the insect meal diet; Segment (+), higher relative abundance in the distal intestine; *foxp3* (+) / *plin2* (+), positive correlation between microbial clade relative abundance and gene expression levels. (b) Representative taxa showing higher relative abundances in salmon fed the reference diet. (c) Representative taxa showing higher relative abundances in salmon fed the insect meal diet. The relative abundances of representative taxa in the feeds are shown as grey dots in panels b and c. As the number of taxa showing significant associations with diet was too high to be properly displayed on the heatmap, we filtered the results to keep those with a q-value < 0.0001. Complete results are available in our accompanying R Markdown reports. Taxa not assigned at the genus level are prepended with letters indicating whether the taxonomic assignment was made at the phylum(p_), order (o_), or family (f_) level. Abbreviations: REF, reference diet; IM, insect meal diet; PI, proximal intestine; DI, distal intestine; FDR, false discovery rate; N.not.zero, number of observations that are not zero.

**Fig. 7.**
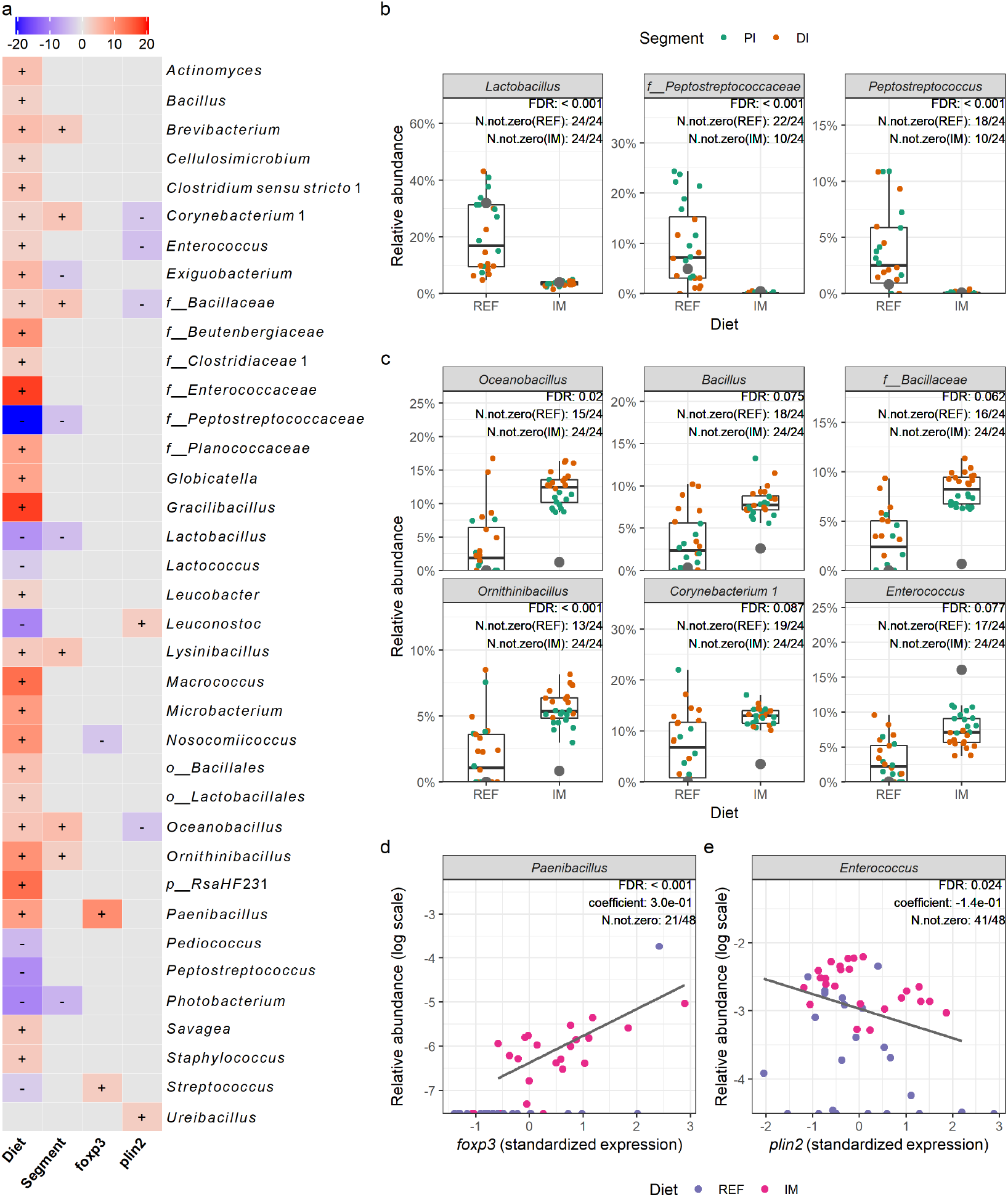
Significant associations between sample metadata and microbial clades in the mucosa. (a) Heatmap summarizing significant associations between sample metadata and microbial clades in the mucosa. Color key: -log(*q*-value) * sign(coefficient). Cells that denote significant associations are colored in red or blue and overlaid with a plus (+) or minus (-) sign that indicates the direction of association: Diet (+), higher relative abundance in salmon fed the insect meal diet; Segment (+), higher relative abundance in the distal intestine; *foxp3* (+) / *plin2* (+), positive correlation between microbial clade relative abundance and gene expression levels. (b) Representative taxa showing higher relative abundances in salmon fed the reference diet. (c) Representative taxa showing higher relative abundances in salmon fed the insect meal diet. (d) Positive correlation between the relative abundance of *Paenibacillus* and *foxp3* expression levels in the intestine. (e) Negative correlation between the relative abundance of *Enterococcus* and *plin2* expression levels in the intestine. The relative abundances of representative taxa in the feeds are shown as grey dots in panels b and c. Taxa not assigned at the genus level are prepended with letters indicating whether the taxonomic assignment was made at the phylum(p_), order (o_), or family (f_) level. Abbreviations: REF, reference diet; IM, insect meal diet; PI, proximal intestine; DI, distal intestine; FDR, false discovery rate; N.not.zero, number of observations that are not zero.

## Discussion

We found that the insect meal diet markedly modulated the Atlantic salmon intestinal microbiota. A group of bacterial genera, dominated by members of the *Bacillaceae* family, was enriched in salmon fed the insect meal diet. These results confirm our previous findings in a seawater feeding trial. We also found that microbiota in the intestine closely resembled that of the feeds. Notably, bacterial genera associated with the diet effects were present in the feeds as well. We conclude that salmon fed the insect meal diets show consistent changes in the intestinal microbiota. The next challenge is to evaluate the extent to which these alterations are attributable to feed microbiota and dietary nutrients.

### Insect meal diet markedly modulated the intestinal microbiota

Higher microbial diversity has been reported in intestinal digesta, and mucosa of salmonids fed diets containing black soldier fly larvae meal (38,61–63). In the present study, however, this was the case for the mucosa, but the opposite was the result for the digesta. Our observation that a particular group of bacterial genera, dominated by members of the *Bacillaceae* family, was enriched in salmon fed the insect meal diet is in line with findings in our previous seawater trial, wherein salmon were fed an insect meal diet containing 15% black soldier fly larvae meal for 16 weeks (38). Among these bacterial genera, *Actinomyces, Bacillus, Brevibacterium, Corynebacterium 1, Enterococcus, Oceanobacillus*, and *Paenibacillus* were also reported to be enriched in rainbow trout fed diets containing 15% or 30% black soldier fly larvae meal (62–64). Similar observations have been made in Siberian sturgeon (*Acipenser baerii*) fed a diet containing 15% BSF larvae meal, inducing higher ab-solute abundances of *Bacillus* and *Enterococcus* (65). In this latter study, fluorescence in situ hybridization (*FISH*) technique was used for the bacteria quantification.

Feed microbiota and dietary nutrients may explain the observed diet effects. We found evidence for the former, because bacterial genera associated with the diet effects were present in the feed samples. Given the hydrothermal treatments during the extrusion step in the feed production, the viability of feed-associated microbes is expected to be low.

As sequencing-based methods cannot differentiate between active (living) and inactive (dormant/dead) microbes, additional work will be needed to elucidate the extent to which the observed diet effects are attributable to the carry-over of inactive microbes and colonization of active microbes from feeds. Methods like viability PCR and RNA sequencing can be applied for such experiments (66). Changes in the feed components may have also contributed to the observed diet effects. For instance, dietary inclusion of soy proteins was suggested to associate with increased relative abundance of lactic acid bacteria in the salmon intestine (29). Thus, the replacement of soy protein concentrate with insect meal may explain the reduction in lactic acid bacteria in salmon fed the insect meal diet. On the other hand, nutrients from the insect meal, such as chitin, may have also promoted the growth of certain bacterial taxa including *Actinomyces* and *Bacillus*. *Actino-myces* species are often identified as active chitin degraders, showing enhanced growth and activity upon chitin addition (67). Many *Bacillus* species are well-known as chitin degraders (68). *Bacillus* was one of the predominant taxa in the intestinal mucosa of salmon fed a chitin-supplemented diet, displaying the highest *in vitro* chitinase activity (69). The latter hypothesis can be tested by supplementing insect meal-specific nutrients to the same basal diet and sequencing the intestinal microbiota of salmon fed these diets.

### Microbiota was similar between intestinal segments

Like its mammalian counterparts (70,71), the salmon intestinal microbiota is also spatially heterogeneous in its composition (72). Specifically, microbial communities differ along the intestinal tract and vary substantially between digesta and mucosa within the same intestinal segment. Due to the batch effects between sequencing runs, we could not directly compare microbial communities in the digesta and mucosa. Nonetheless, our study suggests that conclusions on the diet effect can be different when evaluated using digesta or mucosa samples alone. This is supported by our results showing that diet effects on the alpha-diversity and differential abundance testing were quite different when evaluated independently using digesta or mucosa samples. In contrast, our comparative analysis showed that microbiota variations between intestinal segments were minor in the digesta and neglectable in the mucosa. The diet effects were essentially the same when evaluated using samples from different intestinal segments. Taken together, these results suggest that it may be sufficient to collect digesta and mucosa samples from the distal intestine when conducting a diet-microbiota study in fish with a limited sample size.

### Microbial overlap was low between the intestine and water but high between the intestine and feeds

Water and feed are considered two environmental sources of microbiota which can be transferred to the fish intestine. In line with previous studies in salmon (73–75) and other fish species (76–78), we found that microbial overlap between the intestine and water was low in the present study of salmon in freshwater. This may be explained by the fact that during their freshwater stage, salmon drink little water to accommodate osmoregulation needs in a hypo-osmotic environment, which greatly limits the intake of microbes from the surrounding water environment. Conversely, we found a high overlap between microbiota in the intestine and the feeds. Contradicting results have been reported in the literature regarding microbial overlaps between the fish intestine and formulated feeds (79–82). As discussed earlier, the feed microbiota detected by amplicon sequencing may have primarily originated from inactive microbes. Therefore, feed microbiota can be a confounding factor of the observed diet effects. Given that the influence of feed microbiota on the observed diet effects is unequal across experimental groups as opposed to the water microbiota, we strongly recommend collecting feed samples when designing a sequencing-based, diet-microbiota study in fish.

### Associations between microbial clades and host gene expressions

The close relationship between microbiota and the intestinal immune system is well established (83). Interaction between microbiota and lipid metabolism in the intestine has also been documented (84,85). Here we found a positive correlation between the *Paenibacillus* relative abundance and *foxp3* expression level in the intestine, suggesting a putative link between the enrichment of *Paenibacillus* and increased expression of *foxp3* in salmon fed the insect meal diet. In addition, we found negative correlations between relative abundances of unclassified *Bacillaceae*, *Corynebacterium 1*, *Enterococcus*, *Oceanobacillus*, and unclassified *RsaHF231*, and the expression level of *plin2* in the intestine. This may suggest that the reduction in steatosis in the proximal intestine of salmon fed the insect meal diet might be related to the enrichment of these taxa, either a cause or consequence. However, as microbiome data are sparse and noisy, association analysis is more meaningful when the sample size is much larger than it in this study. Given the limited sample size, our results should be interpreted as exploratory. Further research is required to test if these bacteria taxa are indeed involved in the immune modulation and lipid metabolism in the salmon intestine.

## Conclusion

Our work showed that the insect meal diet markedly modulated the Atlantic salmon intestinal microbiota. Overall, the microbial diversity was lower in the digesta of salmon fed the insect meal diet but higher in the mucosa. A group of bacterial genera, dominated by members of the *Bacillaceae* family, was enriched in salmon fed the insect meal diet. These results support our previous findings from a study of Atlantic salmon in seawater. We also found that microbiota in the intestine closely resembled that of the feed but was distinct from the water microbiota. Notably, bacterial genera associated with the diet effects were present in the feed samples as well. We conclude that salmon fed the insect meal diets show consistent changes in the intestinal microbiota. The next challenge is to evaluate the extent to which these alterations are attributable to feed microbiota and dietary nutrients.

## Data and code availability

The raw 16S rRNA gene sequencing data are deposited at the NCBI SRA database under the BioProject PRJNA730696. The code for reproducing our results is available at the GitHub repository (https://github.com/yanxianl/Li_AqFl1-Microbiota_2021).

## Acknowledgments

Y.L. was granted a scholarship from the China Scholarship Council to pursue his Ph.D. degree at the Norwegian University of Life Sciences. This work was a spin-off of the “AquaFly” project (grant number, 238997), funded by the Research Council of Norway and managed by the Institute of Marine Research. Costs related to this study were covered by the Norwegian University of Life Sciences. The funding agencies had no role in study design, data collection, and interpretation, decision to publish, or preparation of the manuscript.

The authors wish to thank Ellen K. Hage for organizing the sampling and technicians at the Cargill AquaNutrition (former EWOS Innovation) experimental facility for their committed animal care and supports during the sampling.

T.M.K. and Å.K. conceived and designed the study. Y.L., K.G., and Å.K. participated in the sample collection. Y.L., K.G. and A.J.-T. carried out the laboratory works. Y.L. performed the data analysis and completed the first draft of the manuscript. All the authors read, revised, and approved the final version of the manuscript.

We declare no conflicts of interest.

## Supplementary Note 1: Supplemental tables

• Table S1. Contaminating ASVs removed from the feature table (available at the GitHub repository: https://github.com/yanxianl/Li_AqFl1-Microbiota_2021/tree/master/result/table).

## Supplementary Note 2: Supplemental figures

**Fig. S1.**
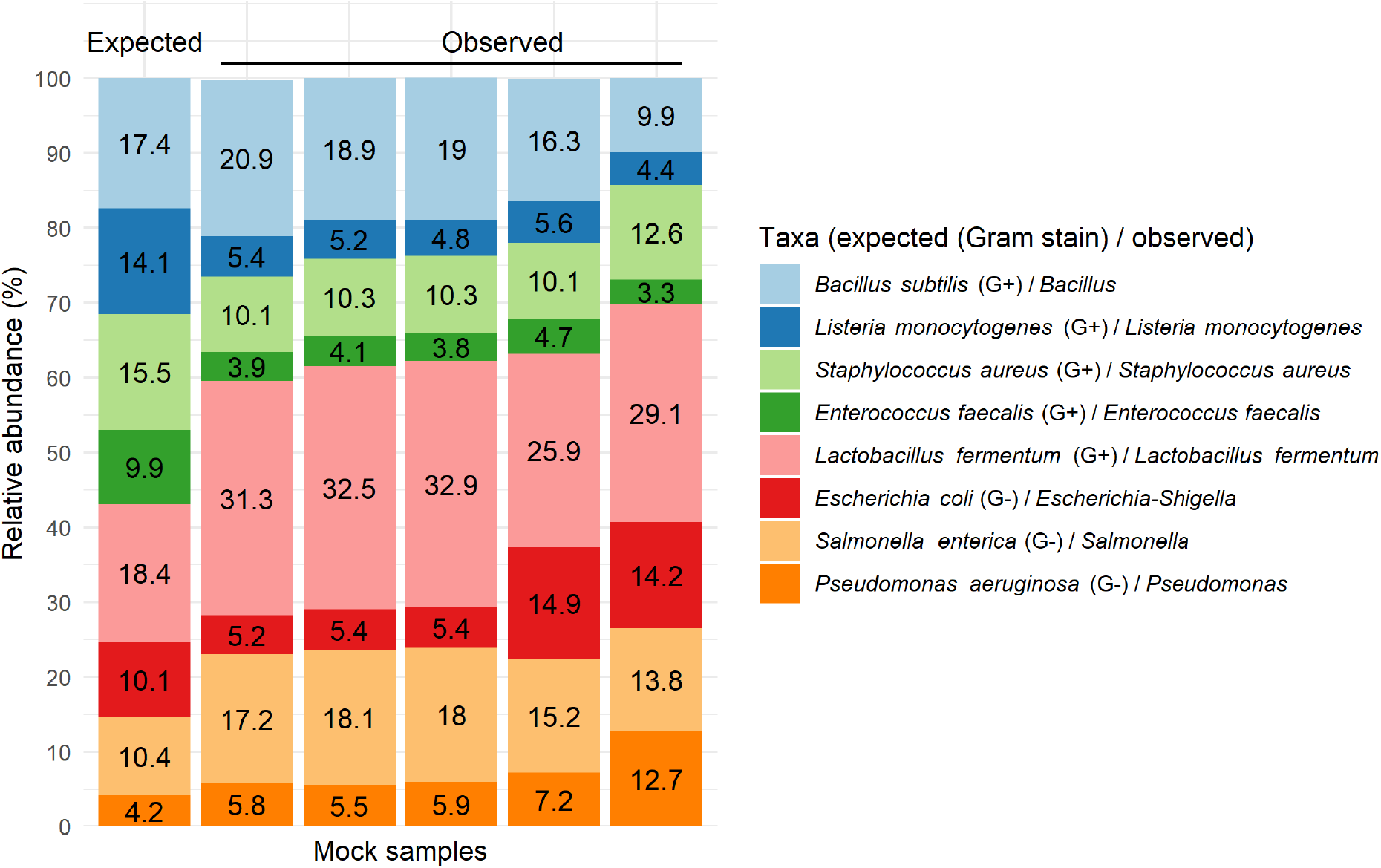
The expected and observed taxonomic composition of the sequenced ZymoBIOMICS mock samples.

**Fig. S2.**
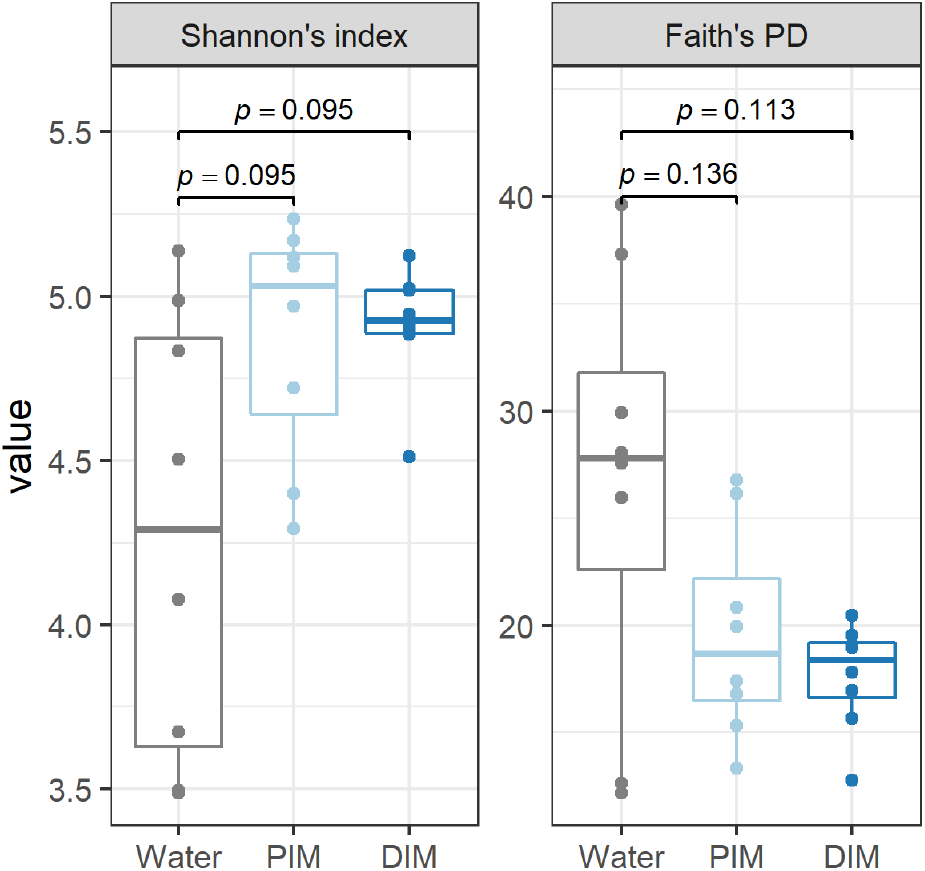
Comparison of alpha-diversity between paired water and intestinal mucosa samples. Note that alpha-diversity indices of intestinal mucosa samples from the same tank were aggregated before running paired t-test. Abbreviations: PIM, proximal intestine mucosa; DIM, distal intestine mucosa; PD, phylogenetic diversity.

**Fig. S3.**
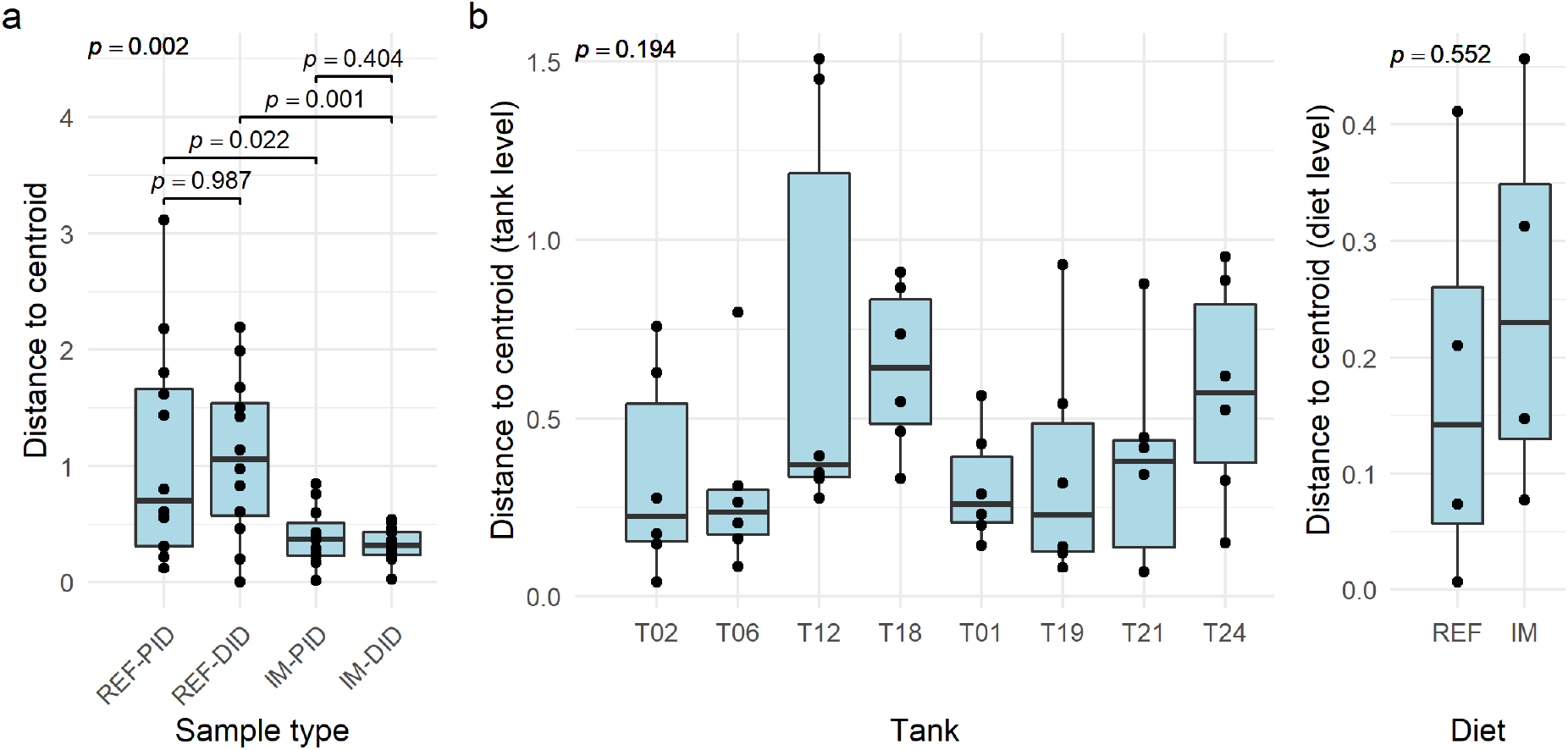
Tests for homogeneity of multivariate dispersions (PERMDISP) in digesta (a) and mucosa (b) samples. (a) The PERMANOVA suggested little evidence of tank effect for digesta samples, thus we used individual fish as the statistical unit when running the PERDISP. (b) The PERDISP, however, was carried out on tank and diet level for mucosa samples because of a significant tank effect. Abbreviations: REF, reference diet; IM, insect meal diet; PID, proximal intestine digesta; DID, distal intestine digesta.

## Notes

### Competing Interest Statement

The authors have declared no competing interest.

